# Tumor-suppressor signature for robust prognosis and prediction in adenocarcinoma non–small cell lung cancer

**DOI:** 10.64898/2026.03.17.712300

**Authors:** Man Jiang, Yuan-De Tan

## Abstract

Non-small cell lung cancer (NSCLC) remains the leading cause of cancer-related mortality, partly due to limited early detection strategies and incomplete understanding of tumor-suppressive mechanisms. In our previous work, we identified 26 tumor suppressor (TS) genes and characterized their biological functions and regulatory networks. We systematically evaluated these TS genes across multiple microarray datasets by analyzing differential expression patterns, correlations with oncogenes, tumor-associated genes, and PD-1–related immune genes, as well as somatic mutation frequencies. A weighted scoring algorithm was used to construct a TS gene signature. Patients were stratified using z-score normalization, and univariable and multivariable Cox proportional hazards models were applied across multiple adenocarcinoma (ADC) cohorts. Prognostic performance was assessed using Kaplan–Meier analysis and AUC metrics. The 26 TS genes were consistently down-regulated in tumors and showed strong negative correlations with oncogenes, particularly in advanced stages. TS genes also exhibited stage-dependent correlations with PD-1–associated immune genes, with chemotaxis/cytokine-signaling genes behaving TS-like, while PDCD1 and SIT1 showed oncogene-like patterns. Somatic mutations were detected in only 32% of LUAD samples for TS genes, compared with 68% for oncogenes. Across seven independent ADC cohorts, high TS-signature expression was associated with significantly reduced risk of death and recurrence/relapse. The TS signature outperformed several published prognostic signatures and demonstrated robust predictive accuracy, with AUC values exceeding 0.7 in multiple datasets and >0.8 for relapse prediction in GSE30219. Across seven independent cohorts, high TS signature expression was consistently associated with significantly reduced risk of death or recurrence/relapse in ADC patients. Patients with high TS signature expression exhibited markedly improved survival probabilities compared with those with low expression. When benchmarked against several established prognostic signatures using AUC metrics, our TS signature demonstrated superior robustness and predictive accuracy for ADC prognosis.

## 1. Introduction

Lung cancer is a leading cause of death worldwide[1], accounting for approximately 20% of deaths caused by cancer[2], and has a high risk of recurrence[3]. The high mortality among patients with lung cancer is mainly due to the absence of an effective screening strategy to identify lung cancer in the early stage[2]. Thus, only ∼25% of patients presenting with lung cancer are in a sufficiently early stage to be amenable to effective surgical treatment [4]. Even after apparent complete resection of NSCLC, 33% of patients with pathologic stage 1A disease and 77% of patients with stage 3A disease die within 5 years of diagnosis[5]. On the basis of histopathological analysis, lung cancer can be divided into two histological cell types: small cell lung cancer (SCLC) and non-small cell lung cancer (NSCLC). NSCLC consists of three subtypes, namely, squamous cell carcinoma (SCC), adenocarcinoma (ADC), and large cell carcinoma (LCC), and accounts for approximately 80% of lung cancer cases [6,7]. Approximately 25∼30% of stage I patients received surgical intervention alone. Despite undergoing curative surgery, more than 25% of patients with stage 1 NSCLC patients die within five years [8]. The high mortality rate of NSCLC patients may be due to a lack of early precision detection, unclear molecular mechanisms, and effective therapeutic methods. The selection of gene signatures and molecular pathways critical for the development of NSCLC could lead to improved therapy [8]. Recently, several study groups have developed gene expression signatures aimed at classifying lung cancer patients into groups with distinct clinical outcomes[8,9,10,11,12,13,14,15,16]. However, most of these gene signatures derived from NSCLC patients are inconsistent across studies. This is because these signature genes may provide discordant information for detection or diagnosis and prognosis because they are up-regulated or down-regulated in different tumor stages and in different cohorts[17]. However, as biomarkers, tumor suppressor (TS) genes do not have these issues because, in normal cells, TS genes are normally expressed and produce special proteins that provide a stop signal that suppresses cell division, slows the cell cycle and marks cells for apoptosis. When TS genes are down-regulated or repressed, the proteins triggering the essential signal that stops the cell division process are not produced, and then, the cells become cancerous[17,18]. For this reason, we selected 26 tumor suppressor genes to construct a gene signature for predicting the survival outcome of NSCLC ADC patients via clinical datasets. In the previous study[17], these 26 tumor suppressor genes were defined by gene function, functional network, pathway, and gene enrichment. Our current study continue to characterize TS-signature by using differential expressions in multiple microarray datasets, correlations with oncogenes, tumor genes, and anti-PD1 genes and somatic mutation profiles and used Cox survival analysis to demonstrate that the TS signature has high prediction in prognosis of NSCLC. Unlike general genes, TS genes are always repressed in cancer but always highly expressed in normal tissues. Hence, the TS signature is more repressed in cancer but higher expressed in normal tissues because expression status of a TS signature are determined by expression status of most TS genes that are accordant and cannot be changed by one or few TS genes that may not be tumor suppressor in other cohorts. We used three clinical cohorts to train the TS signature of 26 TS genes with respect to prognostic and survival outcomes and used 4 cohorts to test this TS signature.

## 2. Materials and Methods

### 2.1. Data collection

A microarray dataset from a cohort of NSCLC patients with ADC was generated by using the GeneChip Human Genome U133 Plus 2.0 expression array [19] and downloaded from the National Institutes of Health (NIH) Gene Expression Omnibus (GEO) website http://www.ncbi.nlm.nih.gov/geo/with <u>access number GSE3141.</u>The data consisted of 54,675 probes, 52 ADC patients and 59 SCC patients. To avoid discordant or confounding outcomes resulting from multiple cell types, we chose microarray data of 52 ADC patients for survival analysis because ADC is the largest cell types in lung cancer. To validate the prediction of overall survival via use of these 26 TS genes (Supplementary **Table S1**) as a signature, we downloaded the other 6 microarray datasets from GEO with access numbers <u>GSE8894</u>[11], <u>GSE68465</u>[13], <u>GSE30219</u>[20], <u>GSE50081</u>[21], <u>GSE37745</u>[22], and <u>GSE31210</u>[23]. GSE8894 was derived from a South Korean cohort consisting of 138 patients collected from 1995-2003, with a median of 44 months of follow-up, of which 63 patients were adenocarcinoma. GSE68465 is a microarray and clinical data from white cohort composed of 443 ADC patients collected from 0.3 months to 98 months in Rockville city. GSE30219 was generated from a French cohort composed of 307 NSCLC patients, among which 85 patients were adenocarcinoma. This cohort has two clinical datasets of death and relapse statuses. GSE50081 is a UHN181 cohort data in which 181 stage-1-2 patients were collected from the University Health Network (UHN) from 1996–2005, however, only 129 patients were available for survival analysis. GSE37745 is a single institute cohort data comprising 196 NSCLC patients whose clinical information and long-term follow-up data were available, of which 106 patients were identified as ADC type. The data also included clinical death and recurrence rates. GSE31210[23] is a cohort data consisting of 226 NSCLC ADC patients subjected to expression profiling selected from 393 stage I-II patients who underwent potential curative resection between 1998 and 2008 at the National Cancer Center Hospital. These data were generated from the Affymetrix microarray platform PGL570, and hence have the same probeid and gene annotation files. These 7 microarray datasets were selected on the basis of the following criteria: (1) adenocarcinoma type, (2) clinical data (death status, survival months, age, sex), (3) non-small cell lung cancer (NSCLC), (4) covariates (such as smoking status, tumor stage, and TNM stage) and (5) were not log-transformation because log-transformed data are not available for standard normalization(z-score). All these 7 microarray datasets selected were generated from cancer patients in single-laboratories and from single-microarray platforms and they did not have technical replicates. Therefore there is not batch effect in these datasets. In addition, these 7 microarray datasets were normalized by authors and used to select genes and to perform survival analysis. So in our study, we were not required to do quality control of these datasets. The basic information (such as sex, age, death/recurrence status, and tumor stage and sample size) and characteristics of these 7 clinical and microarray datasets can be found in Supplementary **Table S2** and more specific detail information of these microarray datasets can be found in authors’ papers that were cited and listed in references.

### 2.2. Ethical Considerations

As this study utilized publicly available, anonymized patient data from the GEO database, which operates under established ethical guidelines, no additional ethical reviews or patient consents were required beyond those obtained by the original investigators. A data availability statement is provided at the end of the Methods section.

### 2.3. Differential expression and correlation analyses

To characterize these 26 tumor suppressor genes, we also downloaded microarray datasets <u>GSE19804</u>[1], <u>GSE103512</u>[24], and <u>GSE40275</u>[25] from GEO of NCBI database. Microarray data GSE19804 were derived from ADC non-smoking female lung cancer in Taiwan and has 15 patients in stage 1A, 19 patients in stage 1B, 12 in stage 2 and 12 in stage 3. Normal samples were derived from adjacent normal lung tissue specimens, that is, tumor and normal samples are paired, so they have the same sample sizes of 58 patients. GSE103512 was made from 280 formalin-fixed and paraffin embedded normal and tumor samples of four cancer types of which 65 patients were breast cancer matched with 10 normal tissues, 57 patients were colorectal cancer matched with 12 normal tissues, 60 were prostate cancer matched with 7 normal tissues. In our current study, 60 non-small cell lung cancer patients matched with 9 normal samples were chosen for showing differential expressions of 26 tumor suppressor genes. GSE40275 was made for differences in gene expression profiles regarding the expression of genes encoding for proteins with G protein-coupled receptor (GPCR) activity between SCLC and NSCLC and normal lung samples where 8 SCLC samples, 16 NSCLC samples and 14 normal lung RNA samples (human) had been purchased from OriGene Technologies. In the present study, we chose 16 NSCLC samples and 9 normal samples for showing differential expressions of 26 tumor suppressor genes. Gene expressions of 26 TS genes between normal and tumor samples were performed using R code (https://github.com/Yuande/run_signature_analysis/) on Affymetrix microarrays <u>GSE19804</u>[1], <u>GSE103512</u>[24], and <u>GSE40275</u>[25] and the results was visualized by pheatmap([pheatmap: Pretty Heatmaps](https://cran.r-project.org/web/packages/pheatmap/pheatmap.pdf)). We also performed correlation analysis of 26 TS genes with 26 oncogenes and genes associated with PD-1 on these three microarray datasets and also used heatmap to visualize these correlation matrices. Heatmap was made by using our R code (https://github.com/Yuande/run_signature_analysis/).

### 2.4. Mutation profiles between the 26 TS genes and 26 oncogenes

To show that our tumor suppressor genes are really different from oncogenes in somatic mutations, we downloaded 568 maf files (568 samples) of LUAD from TCGA and used oncoplot in R pckage maftools to display somatic mutation profiles of 26 TS genes and 26 oncogenes.

### 2.5. Signature scores

We here developed a weight approach to combine the 26 TS genes into one signature for predicting the prognosis of NSCLC patients. First, we utilized the expression data of the 26 TS genes in a cohort to perform univariable or multivariate individual Cox proportional hazard regression analysis, generated 26 p-values for the 26 hazard risks, transformed the p-values to -log values, and finally constructed a weight vector (*w*_*g*_):

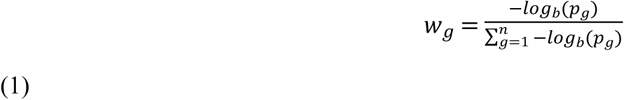

where *n* is number of genes and *g* is the *gth* gene (g = 1, 2, …, n). If gene *g* is a major contributor to survival outcome, then gene *g* has a very small p-value for survival probability, and its weight (*w_g_*) is large; if the gene has a small contribution to survival outcome such that it has a large p-value and small weight (w_g_). Thus, we use the weight vector and gene expression values of genes in patient *i* to construct a new expression value (*x*_*i*_) of the signature (or signature-score) for this patient:

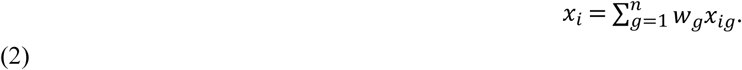

Note that gene expression data and clinical data in a cohort are used to construct a weight set, and the weight set is used to obtain expression values of the signature across patients in the cohort. These signature-scores are utilized to predict the survival outcome of patients. Therefore, our approach does not require training data or validation data. Three cohorts (<u>GSE3141, GSE8894</u>, <u>GSE68465</u>) were used to train signature-score of tumor-signature for Cox hazard proportional survival analysis and 4 cohorts (<u>GSE50081</u>[21], <u>GSE37745</u>, and <u>GSE31210</u>) were used as testing cohorts.

### 2.6. Statistical analysis

Using the expression values of the TS signature in a cohort, we can perform overall and/or recurrence-free survival analysis. To avoid bias or discordant outcomes resulted from multiple NSCLC cell types, only ADC patients were recruited in the survival analysis because ADC type is the largest group in lung cancer. The ADC patients were divided into three groups by means of standard normalized signature expression (z-score) [26]: high-expression group where z-score was > 0.2, low-expression group where z-score was < -0.2, and medium-expression group where -0.2 ≤ z-score ≤ 0.2. Survival analysis was performed to compare the high- and low-expression groups. The medium-expression group is a noise group and discarded. There is no theoretical criterion to select patients for survival analysis, our practice is that if the cutoff is out of ± 0.2, then it will lead to a result that high- and low-expression groups have too less patients, which will result in a serial bias in survival analysis. We found that a z-score < -0.2 and z-score > 0.2 are the best criteria because, in the 7 cohorts, this criterion works well for classification. The z-scores follow standard normal distribution with mean = 0 and variance = 1. The range of ―0.2 ≤ z-score ≤ 0.2 is a noise interval where expression of signature in patients are uncertain and noise due to random effect and hence not correlated with survival status of patients. In the high-expression group with z-score > 0.2, tumor suppressor genes are normally or highly expressed, tumors are repressed so that the patients have good survival status or high alive probability, while in the low-expression group, tumor suppressor genes are repressed, cancer or tumors are growing, and the patients have bad survival status or have high risk probability of death. Multivariate Cox proportional hazard regressions with covariates of cancer stage, patient age, sex and/or smoking status, therapy, treatment adjuvant etc were implemented to construct a predictive survival model. The Wald test was used to test for associations of differential expression of the TS-signature with survival outcome, and Kaplan-Meier curves were generated to visualize survival differences between the high- and low-risk patient groups. Correlation analysis of TS genes with oncogenes and immune genes was conducted in R environment.

### 2.7. Evaluation of Receiver operating characteristic (ROC)

Receiver operating characteristic (ROC) analysis was used to assess the accuracy of unvariate Cox hazard proportional model predictions by plotting sensitivity or true positive rate (TPR) versus 1-specificity or false positive rate (FPR) as the threshold varies over an entire range of diagnostic test results. We performed R packages ROCR and gplots to do ROC analysis of Cox hazard proportional model predictions of ADC patients classified by using gene-signature in the cohorts GSE8894, GSE30219 (death status), GSE3141 or GSE50081 (recurrence and death statuses). After performing ROC analysis, we calculated the **area under ROC curve (AUC)** of each gene-signature. AUC represents the probability that the signature predicts survival of ADC patients over an entire range of diagnostic test results.

### 2.8. R package for signature survival analysis and R code for differential and correlation analyses

An R package SignatureSurival was created and recruited by R CRAM (CRAN -Package signatureSurvival (r-project.org). The package can be used to do screen of signature genes, weight vector and signature score calculations, performances of univariable and multivariate linear Cox proportional hazard survival analyses and making forest plots of results of survival or overall survival analysis of patients using signature. We used R code run_signature_analysis.R (https://github.com/Yuande/run_signature_analysis/) to show differential expressions of TS genes on three microarray datasets and correlation of 26 TS genes with 26 oncogenes and immune genes related with PD1.

## 3. Results

### 3.1. Differential expressions of the 26 TS genes

To show broadly differential expressions of our 26 TS genes between normal and ADC tumor samples, we downloaded microarray datasets GSE19804, GSE103512 [24], and GSE40275[25]. GSE19804 was generated from female non-smoking cohort in Taiwan and contains tumor samples from patients in stage1A, stage1B, stage2 and stage3 and paired normal samples. GSE103512 was generated from institute of pathology/LMU Munich and includes 40 NSCLC ADC patients matched with 9 normal samples. GSE40275 was made from Origene Technologies cohort and has 16 NSCLC ADC samples matched with 9 normal samples. We used heatmap to separately display differential expressions of the 26 TS genes between normal and tumor samples in stage 1A, stage1B, stage2 and stage3. Figure1A exhibits week expression difference between normal and tumor samples in stage 1A. The expression difference became obvious in stage 1B(Fig.1B) and differential expression became strong in stage 2(Fig.1C) and very strong in stage3(Fig.1D). In tumor stages 2-3, almost all normal samples had higher expression than tumor samples. In LMU Munich cohort (GSE103512), the 26 TS genes also show higher expression in 9 normal samples compared to NSCLC ADC cell, K-cells, NSCLC-coex-paths, R-cells and P-cells (Fig.S1), except that TS gene RAP1A had low expression in normal samples P-cells and NSCLC-coex-paths. Compared to Figure 1A, these tumor cells should be in more early stage. In Origene Technologies cohort (GSE40275), most of the 26 TS genes had higher expressions in normal samples than in tumor samples (Fig.S2). TS genes RHOB, SASH1, GPC3, KCNRG, LATS2, EXT1 and TBRG1 did not show differential expressions between normal and tumor samples. This case is similar to that in stage 1A in GSE19804. This can infer that patients in GSE40275 cohort were in the stage 1A.

**Figure 1.**
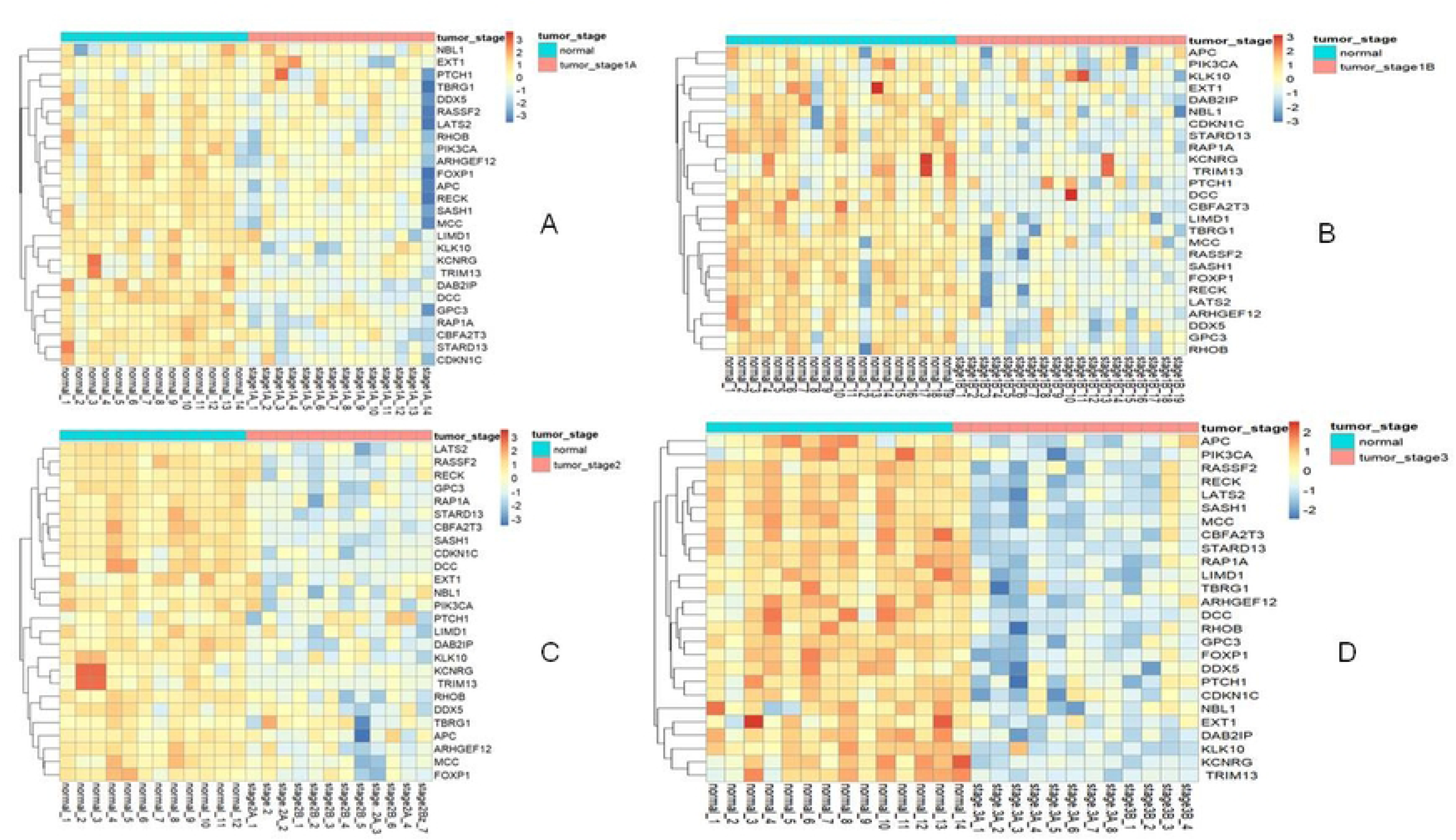
Heatmaps for differential expressions of the 26 TS genes between normal and tumor samples in tumor stages Microarray data was generated by the Affymetrix microarray platform PGL570 from Taiwan female non-smoking cohort. A: stage 1A. B: stage 1B. C: stage 2. D: stage 3.

### 3.2. Correlations of the 26 TS Genes with Oncogenes

To investigate how tumor suppressor (TS) genes interact with oncogenic pathways during tumor progression, we randomly selected 26 oncogenes from a curated oncogene list and examined their expression correlations with our 26 TS genes. Biologically, oncogenes are typically upregulated in tumor tissues, whereas TS genes are repressed; therefore, negative expression correlations between the two groups are expected. Using microarray data from the Taiwan female nonsmoking cohort, we assessed dynamic correlation patterns across tumor stages (stage 1A to stage 3). In stage 1A, most TS genes showed strong positive correlations with ATR, CDK2, NOTCH3, and KDM6A, weak positive or negative correlations with ATM, NF1, RAD51D, TP53, and PHF6, and strong negative correlations with the remaining oncogenes (Fig.2A). In stage 1B, TS genes such as PIK3CA, EXT1, and ARHGEF12 exhibited strong positive correlations with NF1, RAD51D, and KDM6A, and weak positive correlations with several other oncogenes, while most TS genes maintained strong or weak negative correlations with the remaining oncogenes (Fig.2B). In stage 2, the TS genes were strongly negatively correlated with PHF6, IDH1, IDH2, EZH2, DNMT3B, FOXM1, FANCA, CCNE1, and BARC, but strongly positively correlated with CDK2, NF1, NOTCH3, RAD51D, KIT2, and ATM (Fig.2C). By stage 3, the TS genes showed only weak positive or weak negative correlations with NF1, mTOR, NOTCH3, and KDM6A, but remained strongly negatively correlated with most other oncogenes (Fig.2D). The color scale indicates that positive correlation coefficients ranged from 0 to 0.2, whereas negative correlations ranged from −0.2 to −0.8. Thus, the 26 TS genes were predominantly and strongly negatively correlated with oncogenes in stage 3. These findings are consistent with the differential expression patterns of the TS genes shown in Figure 1.

**Figure 2.**
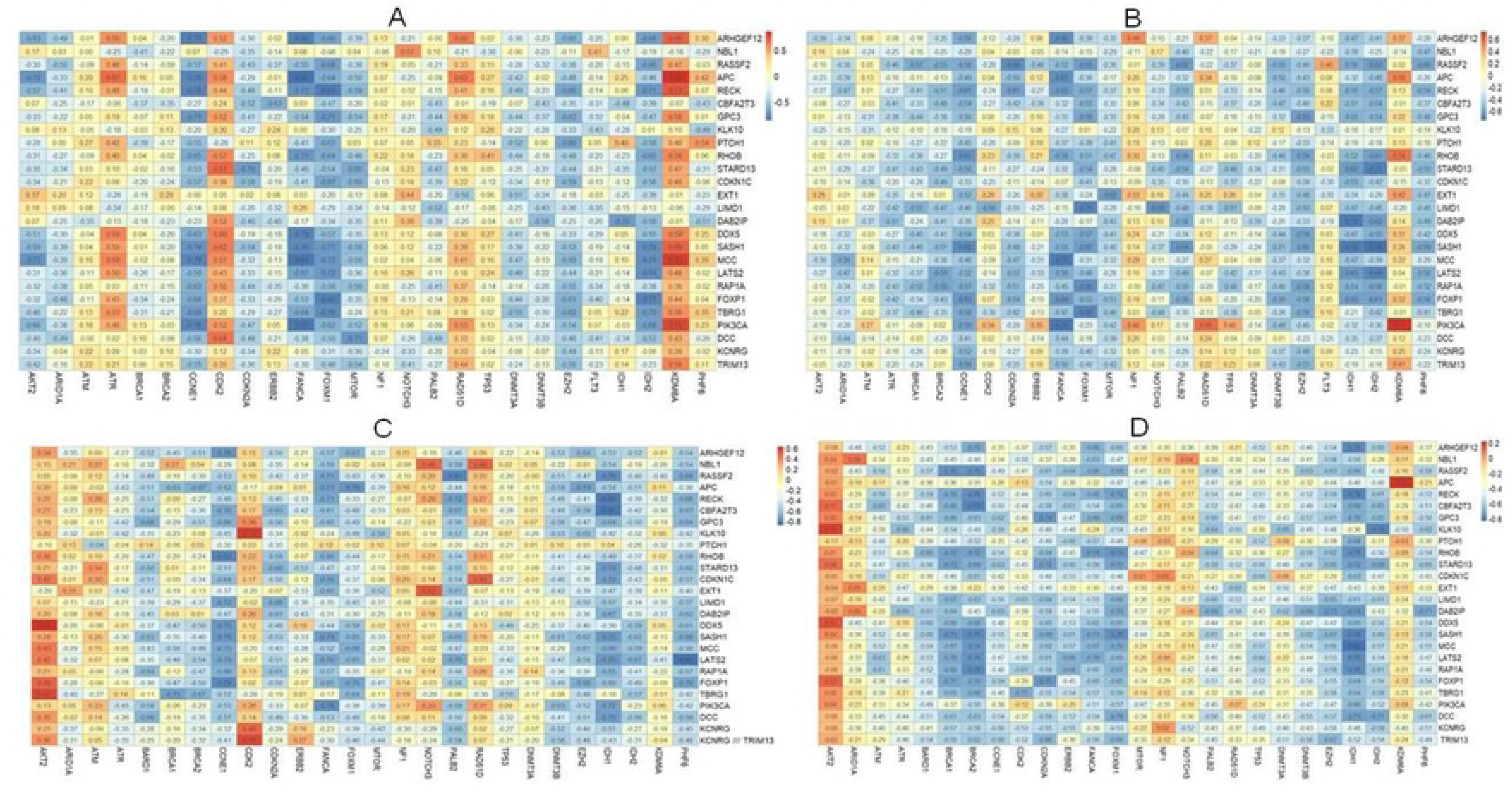
Heatmaps for correlation of the 26 TS genes (in row) with 26 oncogenes (in column) in tumor stages. Microarray data was generated by the Affymetrix microarray platform PGL570 from Taiwan female non-smoking cohort. A: stage 1A. B: stage 1B. C: stage 2. D: stage 3.

### 3.3. Correlations of the 26 TS Genes with Genes Associated with PD-1

To investigate how tumor suppressor (TS) genes interact with immune-related pathways, we examined their expression correlations with genes associated with PD-1 signaling. Mandal et al. [27]identified genes differentially expressed between MSI-H parental cells treated with anti–PD-1 and MSI-H isotype controls. Genes upregulated in anti–PD-1–treated parental cells and downregulated in isotype controls were classified as cluster 1, whereas genes downregulated in anti–PD-1–treated parental cells but upregulated in isotype controls were classified as cluster 2.

Cluster 1 contains host immunity genes, including those involved in lymphocyte activation/function (CD3G, SKAP1, ZAP70, SLA2, SIT1, CD247, LCK, CD2), immune regulation/checkpoint control (FASLG, CTLA4, LAG3, PDCD1), immune cytolytic activity (not detected in GSE19804), and chemotaxis/cytokine signaling (IFNG, CXCR6, CCL4, CXCL9, CCL5, CCL8). Cluster 2 includes genes associated with stromal or tumor biology, grouped into melanin metabolism (CITED1, DCT, GPR3, NYAP1, RGS11, SLC45A2, SOX10, TYR, TYRP1) and tumor metabolism (ALX1, FABP7, PAX3, ZIC2). All of these genes were present in the GSE19804 dataset.

Using GSE19804 microarray data, we evaluated correlations between the 26 TS genes and cluster 1 host immunity genes across tumor stages (Fig.3). The TS genes showed strong or weak positive correlations with the six chemotaxis/cytokine signaling genes (IFNG, CXCR6, CCL4, CXCL9, CCL5, CCL8) in stages 1A, 1B, 2, and 3(Fig.3A-D). This pattern indicates that these chemotaxis genes behave similarly to TS genes—highly expressed in normal tissues and downregulated in tumors.

**Figure 3.**
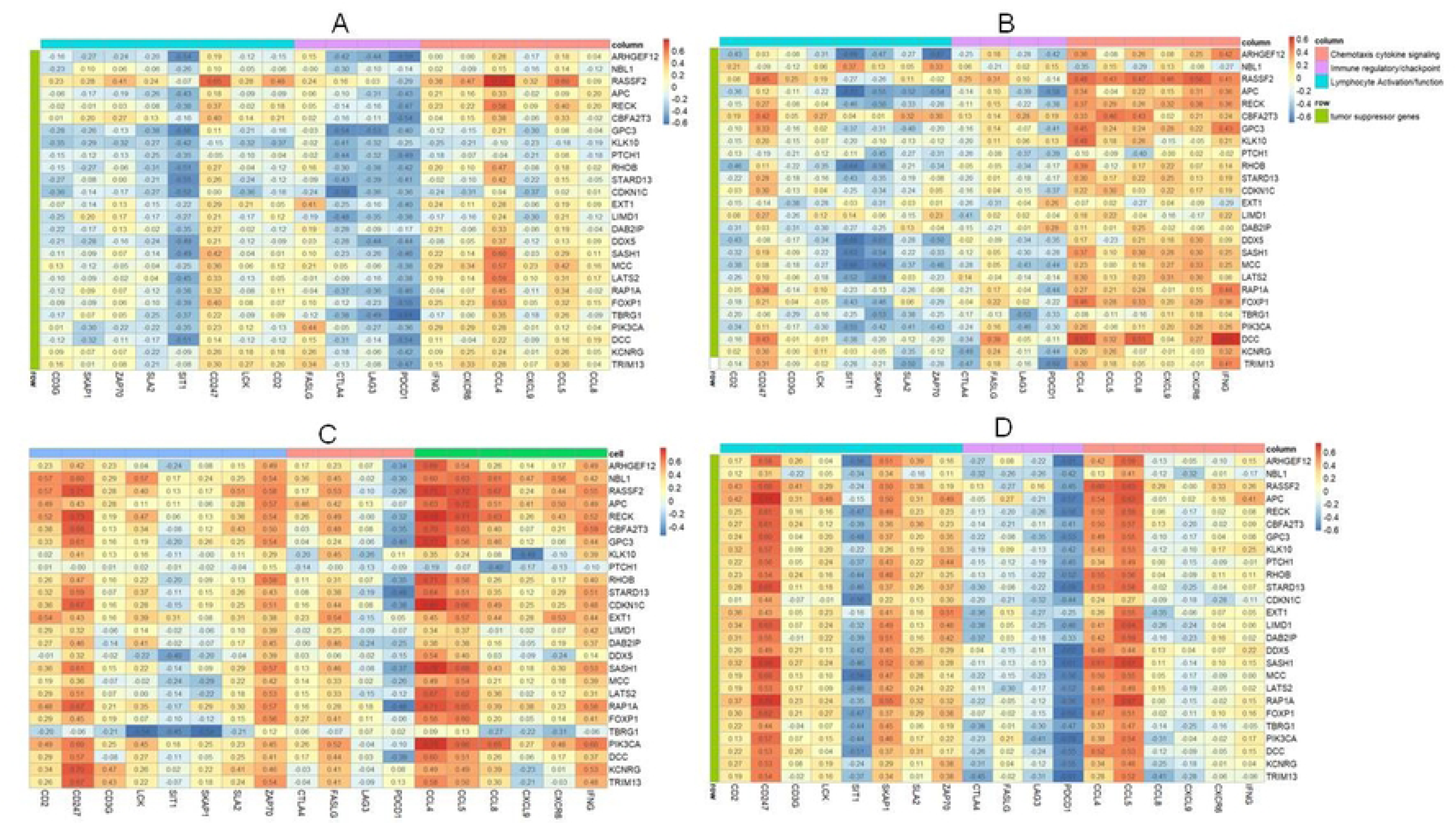
Heatmaps for correlation of the 26 TS genes (in row) with host immunity genes (in column) in tumor stages. Microarray data was generated by the Affymetrix microarray platform PGL570 from Taiwan female non-smoking cohort. A: stage 1A. B: stage 1B. C: stage 2. D: stage 3.

In stage 1A, the TS genes were predominantly negatively correlated with immune regulatory/checkpoint genes and lymphocyte activation/function genes (Fig.3A). By stage 1B, CD247 (lymphocyte activation/function) and FASLG (immune regulatory/checkpoint) shifted to strong positive correlations with nearly all TS genes, while the remaining genes in these categories remained negatively correlated (Fig.3B).

In stage 2, all lymphocyte activation/function genes except SIT1 became strongly positively correlated with the TS genes, whereas all immune regulatory/checkpoint genes remained negatively correlated (Fig.3C). By stage 3, all lymphocyte activation/function genes except SIT1, and all immune regulatory/checkpoint genes except PDCD1, showed strong positive correlations with most TS genes (Fig.3D). These results suggest that as tumors progress, many immune activation and regulatory genes become TS-like, reflecting their reduced expression in tumor tissues.

Two genes—SIT1 and PDCD1—remained strongly negatively correlated with all 26 TS genes across all stages, resembling oncogene-like behavior. PDCD1 (PD-1) encodes an immune-inhibitory receptor expressed on activated T cells and plays a central role in regulating effector CD8+ T-cell function and anti-tumor immunity. PDCD1 is expressed in multiple tumor types, including melanoma, hepatocellular carcinoma, and NSCLC. In the Taiwan female nonsmoking cohort, PDCD1 was consistently downregulated in normal tissues and upregulated in tumor tissues across all stages, consistent with its negative correlation with TS genes.

We next examined correlations between the 26 TS genes and cluster 2 genes (Fig.S3). In early stages (1A and 1B), TS genes showed weak negative correlations with tumor metabolism genes (FABP7, PAX3, ZIC2, ALX1) (Fig.S3A-B). In stage 2, these correlations strengthened, with TS genes showing weak to strong negative correlations with all four tumor metabolism genes (Fig.S3C). By stage 3, TS genes were strongly negatively correlated with FABP7, PAX3, and ZIC2, and weakly negatively correlated with ALX1 (Fig.S3D). These patterns are expected because tumor metabolism genes exhibit expression patterns opposite to TS genes.

Melanin metabolism genes are not directly associated with tumorigenesis. Accordingly, TS genes showed variable correlations with these genes in stages 1A and 1B, ranging from weakly positive to weakly negative or no correlation (Fig.S3A-B). In stages 2 and 3, TS genes were strongly positively correlated with TYR and TYRP1, but strongly negatively correlated with SOX10 and SLC45A2, while showing minimal or weak correlations with the remaining melanin metabolism genes (Fig.S3C-D).

### 3.4. Mutation Profiles of the 26 TS Genes and 26 Oncogenes

To compare the somatic mutation landscapes of the 26 tumor suppressor (TS) genes with those of 26 oncogenes, we merged 568 MAF files from adenocarcinoma non–small cell lung cancer (LUAD) samples downloaded from TCGA. Somatic mutation profiles were visualized using maftools. As shown in Figure 4, the TS genes and oncogenes displayed markedly different mutation patterns.

**Figure 4.**
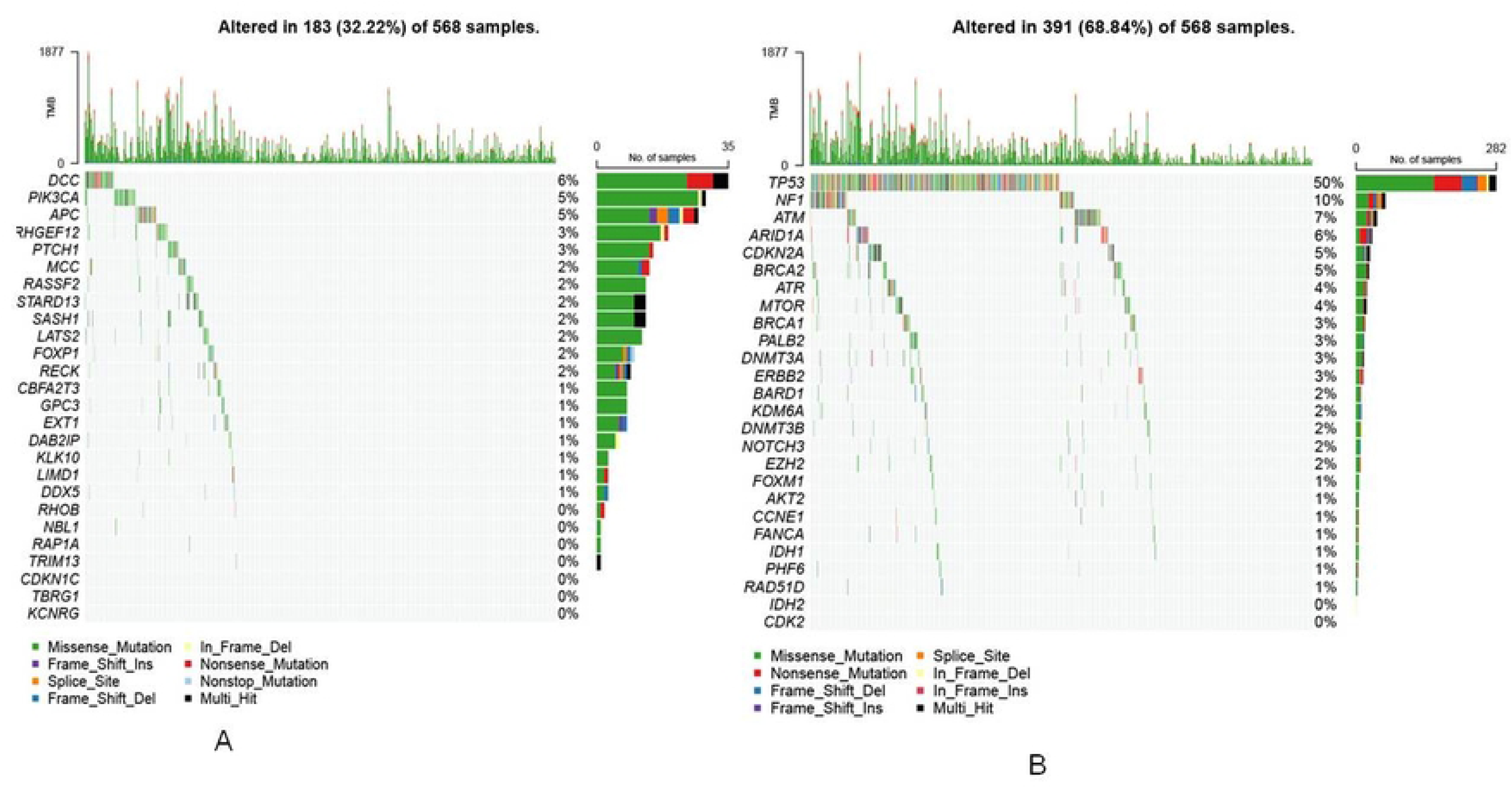
Mutation distributions in 568 LUAD samples. 568 LUAD (NSCLC ADC cancer) maf files were downloaded from TCGA website and merged together by using maftools. The mutation distributions of the 26 TS genes (A) and 26 oncogenes were made by using oncogeneplots.

The 26 TS genes exhibited a relatively low mutation burden, with mutation frequencies ranging from 6% (DCC) to 1% (LMD1). Mutations in TS genes were detected in only 183 LUAD samples (32.22%) (Fig.4A). In contrast, the 26 oncogenes showed two distinct mutation patterns, with mutation frequencies ranging from 50% (TP53) to 1% (RAD51D). Overall, oncogene mutations were present in 391 LUAD samples (68.48%) (Fig.4B).

These results demonstrate that somatic mutations contributing to tumorigenesis are substantially more common in oncogenes than in TS genes. The lower mutation frequency in TS genes suggests that their tumor-suppressive functions may be more frequently disrupted through mechanisms other than direct coding mutations, such as transcriptional repression, epigenetic silencing, or pathway-level dysregulation.

### 3.5. Survival analysis

#### 3.5.1. Survival analysis of ADC patients in death status using TS signature

In this and next sections, we focus on discussing whether our TS genes can robustly be applied to predict prognosis of NSCLC ADC patients using clinical training and testing datasets. To use our TS genes to predict prognosis of cancer patients, we combined the 26 TS genes to a signature using our algorithm. Step 1: we performed Cox proportional hazard survival analysis on three training cohorts to train these 26 TS genes and output 26 p-values in each cohort. Step2: we transformed p-values to –log_2_ values. Step 3: we calculate TS-signature scores across all patients in three training cohorts by using Eq.1 and Eq.2. Step4: standard normalization (z-score) of TS-signature scores was made across all patients. Step 5: patients in a cohort were divided into three groups: high-expression group with z-score > 0.2, low-expression group with z-score < -0.2 and medium-expression group with -0.2 ≤ z-score ≤ 0.2. We discarded medium-expression group and re-performed Cox hazard proportional survival analysis of NSCLC patients on the basis of the clinical death status data. The results obtained from the GSE8894 cohort revealed that the high-expression patients had very significantly higher survival probabilities than the low-expression patients and significantly reduced death risk with p-value = 0.00349 for hazard risk (HR) = -1.1369 in the univariable model (Fig.5a) and p-value = 0.01149 for HR = -1.0403 in the multivariate model adjusted for sex and age (Fig.5d). Age and sex did not impact survival outcome (p-value = 0.9195 for age and 0.9719 for sex, (Fig 10b). In the GSE3141 cohort, the patients with high expression of the TS- signature also had a significantly higher alive probability than those with low expression (p-value = 5e-05 for HR = -2.5838, (Fig.5c).

**Figure 5.**
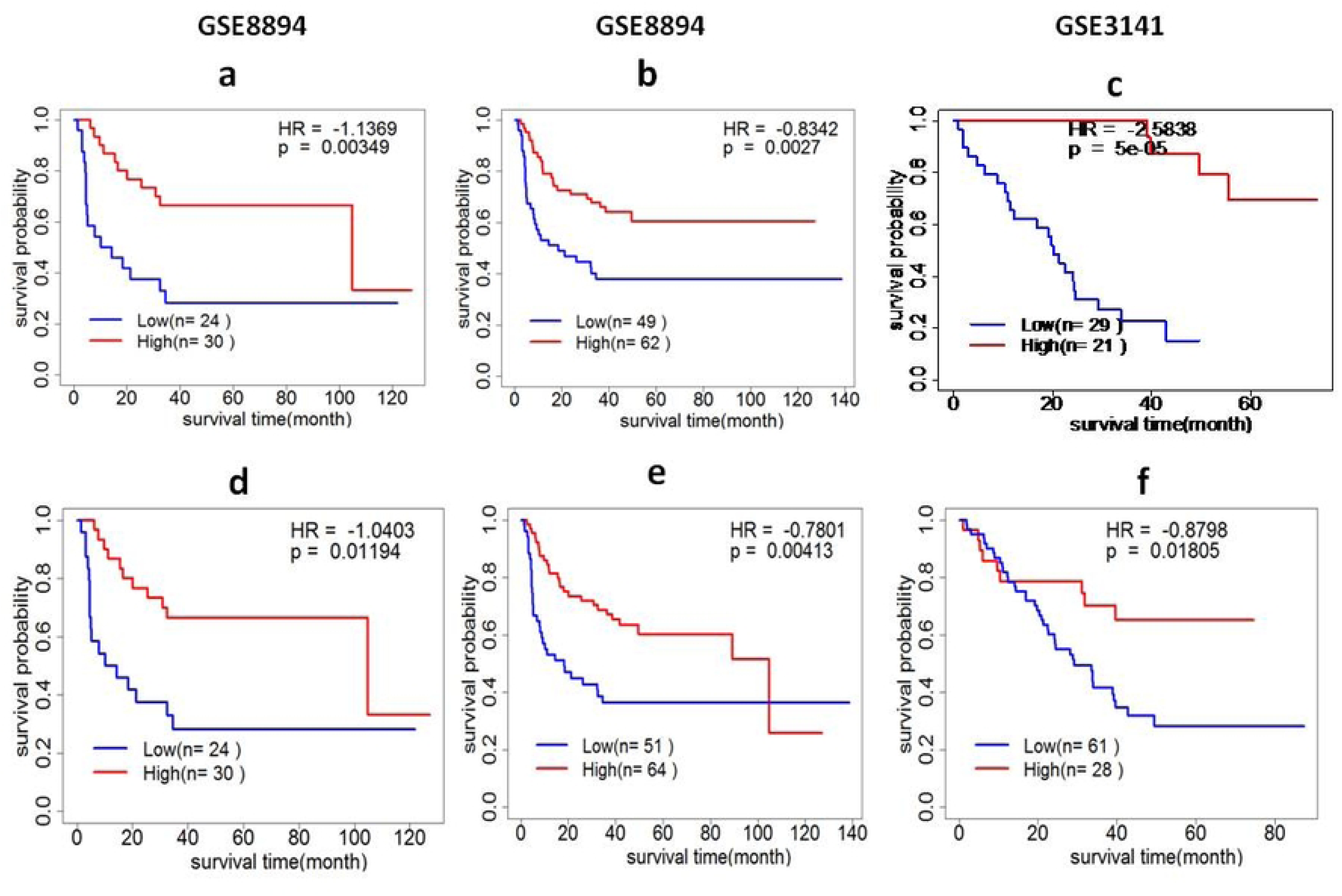
Cox proportional hazard survival analysis of NSCLC patients in the GSE8894 and GSE3141 cohorts. GSE8894 is a cohort composed of 138 lung cancer patients, of which 62 patients had ADC, 75 had squamous cell carcinoma (SCC), and 69 patients died. This cohort had recurrence data. GSE3141 is a cohort composed of 112 patients, of which 58 patients had ADC, 54 had SCC, and 32 died. Overall survival (OS) analysis of ADC (**a**) and ADC/SCC (**b**) patients in the GSE8894 cohort was conducted via a univariable Cox proportional hazard model. OS analysis of ADC (**c**) and ADC/SCC (**f**) patients who died in the GSE3141 cohort was conducted via a univariable Cox proportional hazard model. OS analysis of ADC (**d**) and ADC/SCC (**e**) patients who died was conducted via a multivariate Cox proportional hazard model adjusted for age and sex in the GSE8894 cohort.

In GSE50081 cohort, our survival analysis of the 129 ADC patients was conducted with univariable and multivariate models adjusted for age, sex, tumor stage, and/or smoking status. In the univariable model(Fig.6a) and multivariate models adjusted for age and sex (Fig.6b), adjusted for age, sex, and smoking (Fig.6c), adjusted for age, sex, and tumor stage t (Fig.6d), adjusted for age, sex and tumor stage n (Fig.6e), and adjusted for age, sex, and tumor stage *m (*Fig.6f), survival probability was significantly higher in the high-expression patients than in the low-expression patients with p-value <0.001 for HR< -0.89, meaning that patients with higher expression of TS signature had small death risk. Figure 10e shows that the covariates age, sex, and smoking did not impact the survival outcomes of patients; however, tumor stages *t*, *n*, and *m* significantly increased the risk of death in patients with p-value <0.05.

**Figure 6.**
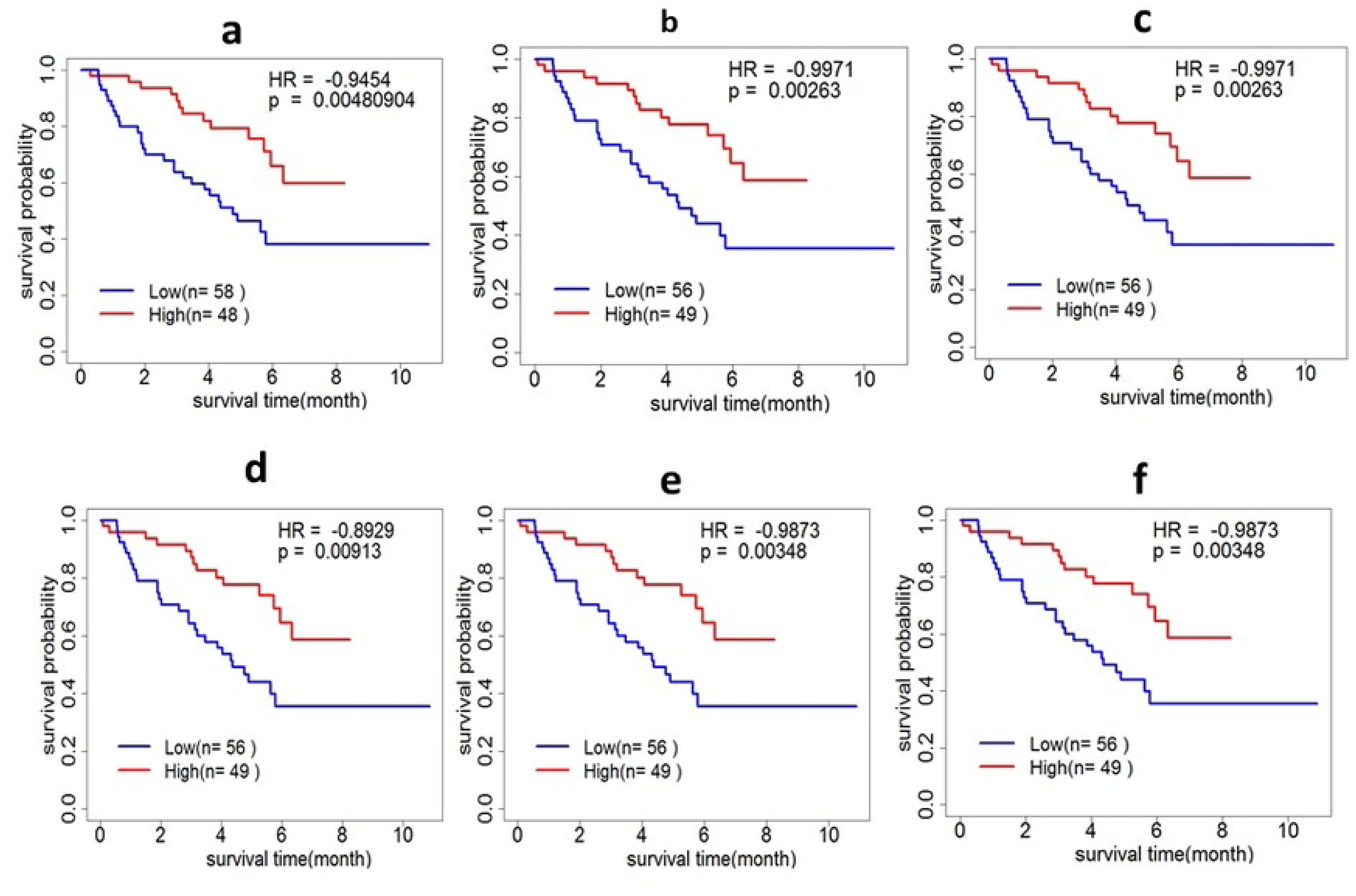
Cox proportional hazard survival analysis of NSCLC patients with all stages of death in the GSE50081 cohort GSE50081 included 181 NSCLC patients, 129 of whom had ADC and 52 of whom had SCC. OS analysis of 129 ADC patients who died was conducted via a univariable Cox proportional hazard model (**a**); multivariate Cox proportional hazard model1 adjusted for age and sex (**b**); multivariate model2 adjusted for age, sex, and smoking (**c**); multivariate model3 adjusted for age, sex, and stage t (tumor size) (**d**); multivariate model4 adjusted for age, sex, and stage n (node) (**e**); and multivariate model5 adjusted for age, sex, and stage m (metastasis) (**f**).

GSE31210 is a testing cohort consisting of 226 ADC patients among which 168 patients were diagnosed in stage-1 and 58 patients in stage-2. Among the 226 ADC patients, 204 underwent complete resection and did not receive postoperative chemotherapy and/or radiotherapy unless they relapsed. Our survival analysis of the 204 ADC patients was conducted via a univariable model (Fig.S4a); multivariate model 1 adjusted for age and sex (Fig.S4b), model 2 adjusted for age, sex, and smoking status (Fig.S4c); model 3 adjusted for age, sex, and genotype (Fig.S4d) where the genotype included ALK-fusion**^+^**, EGFR**^−^**KRAS**^−^**ALK**^−^**, EGFR mutation**^+^**, ALK-fusion**^+^**, and KRAS mutation^+^; univariable model in stage1 (Fig.S4e); model 4 adjusted for age and sex in stage1 (Fig.S4f); model 5 adjusted for age, sex, and smoking status in stage1 (Fig.S4g); and model 6 adjusted for age, sex, and genotype in stage1 (Fig.S4h). The results are summarized in Figure S4. The first four models yielded similar results: HR< -1.75, survival probability was significantly higher in the high-expression patients than in the low-expression patients with p-value < 0.0011 (Fig.S4a-d), Figure S4e-h also displays similar results. Compared with Figure S4a-4d, we found that the alive probability was higher in the stage-1 patients than in the stage-1-2 patients, especially when the censor time was more than 100 months. The results suggested that the patients with high-expressions of the TS signature had less death risk than those with low-expressions of the TS signature. Figure 10c shows the outcomes of multivariate proportional hazard survival analysis in the GSE31210 dataset. In stage1 and stage-1-2 patients, the covariates age, sex, smoking status, and genotype were not significantly associated with survival outcomes in patients with p-values > 0.05 (Fig.10c).

In GSE30219 cohort, 293 patients were diagnosed with lung cancer, among which 85 patients were identified as ADC type. We used the TS signature to classify the 85 ADC patients into three groups: high-expression, low-expression and medium-expression groups. Overall survival analysis was conducted with a univariable model and a multivariate model adjusted for age and sex (not adjust for tumor stage because there was no variation in stages *t* (tumor size), *n* (node), and *m* (metastasis)). The TS signature significantly reduced the death risk of patients with p-value = 0.00223 for HR = -1.4858 in the univariable model (Fig.7a) and with p-value = 0.00419 for HR = -1.6576 in the multivariate model (Fig.7b). Survival probability was higher in the high-expression patients than in the low-expression patients (Fig.7a-b). As shown in Figure 10d, we did not observe that covariate sex was significantly associated with death risk, but age significantly increased the death risk of patients with p-value = 0.025 for HR= 0.0451.

**Figure 7.**
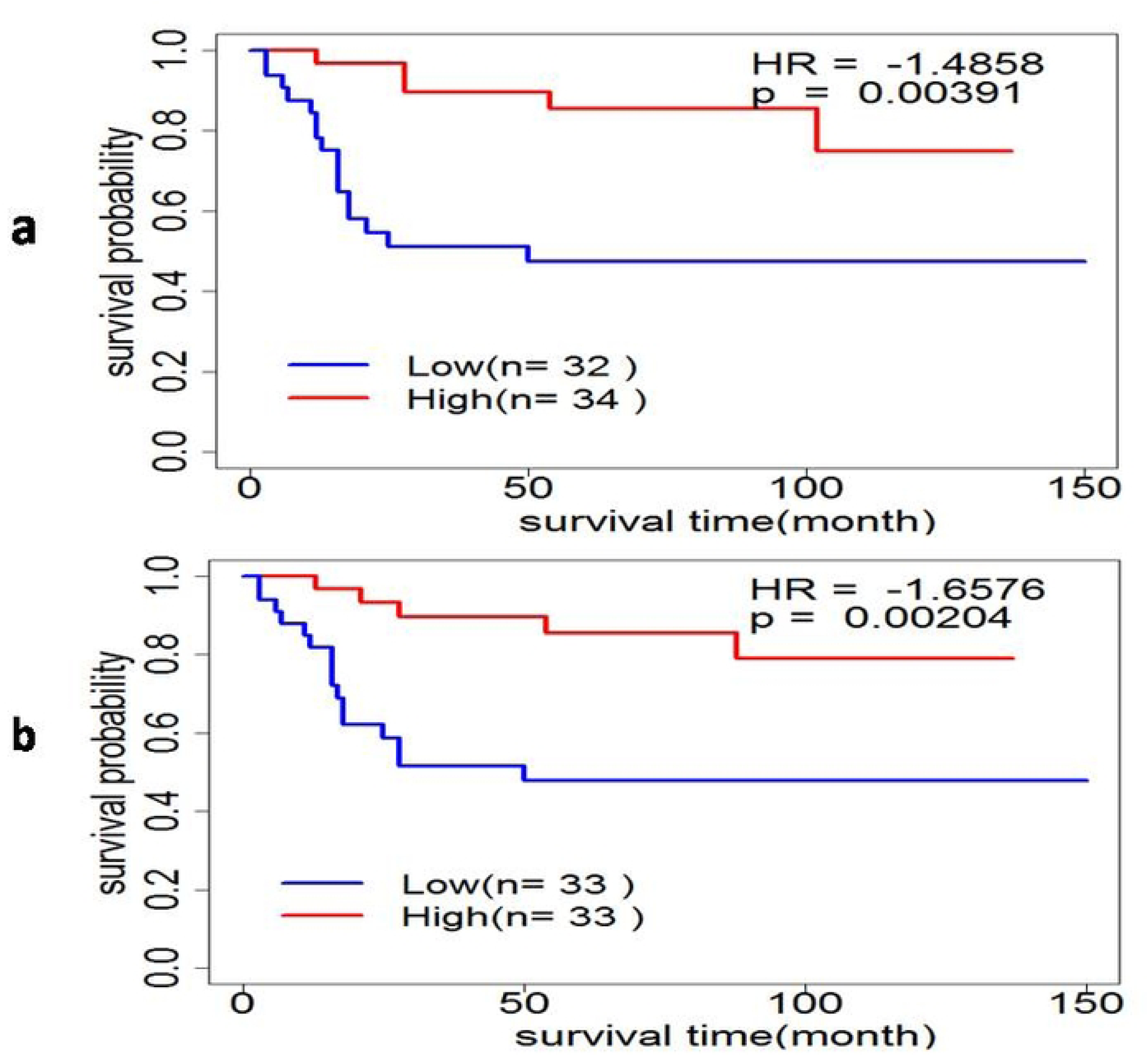
Cox proportional hazard survival analysis of NSCLC patients according to death status in the GSE30219 cohort GSE30219 included data from 293 NSCLC patients, of which 85 patients had ADC, 61 had SCC, and 147 had other types. OS analysis of ADC patients who died was conducted via a univariable Cox proportional hazard model (**a**) and a multivariate Cox proportional hazard model adjusted for age and sex (**b**).

GSE68465 is a Rockville cohort consisting of 442 ADC patients with high-quality gene expression data, pathological data, and clinical information describing the severity of the disease at surgery and the clinical course of the disease after sampling. We performed Cox proportional hazard survival analysis of 442 ADC patients in GSE68465 with univariable and multivariate models adjusted for age, sex, and/or smoking status, chemotherapy, and tumor stages *t* and *n*. The results show that in the univariable model (Fig.S5a), multivariate model 1 adjusted for age and sex (Fig.S5b); model 2 adjusted for age, sex, and smoking status (Fig.S5c); model 3 adjusted for age, sex, and chemotherapy (Fig.S5a), high expression of the TS signature significantly reduced the risk of death with p-value < 0.05 for HR< -0.6908. The high-expression patients had significantly higher survival probabilities than the low-expression patients with p-value <0.05. However, in multivariate models 4 and 5 adjusted for age and sex, tumor stage *t* (Fig.S5e) or tumor stage *n* (Fig.S5f), there was no significant difference in survival probability between the high- and low-expression patients, indicating that tumor stages *t* and *n* did not have effect on the survival outcomes.

In GSE37745, 196 patients were diagnosed with lung cancer, of which 106 patients were adenocarcinoma, 66 were squamous cell carcinoma, and 24 were large cell carcinoma. The overall Cox proportional hazard survival regressions of 106 ADC patients in death status in stage 1 and all stages were conducted with univariable and multivariate models. As seen in the GSE3141, GSE30219, GSE8894, and GSE50081 cohorts, the hazard risk was less than -0.6 for all patients, and the high-expression patients had a significantly higher survival probability than the low-expression patients with p-value = 0.0167 in the univariable model for stage 1 (Fig.8a), 0.0132 in all stages (Fig.8b), 0.0178 in the multivariate model adjusted for age and sex for stage 1 (Fig.8c), and 0.0135 in all stages (Fig.8d). The covariates age and sex in stage1 and all stages were not associated with patient survival (Fig.10a).

**Figure 8.**
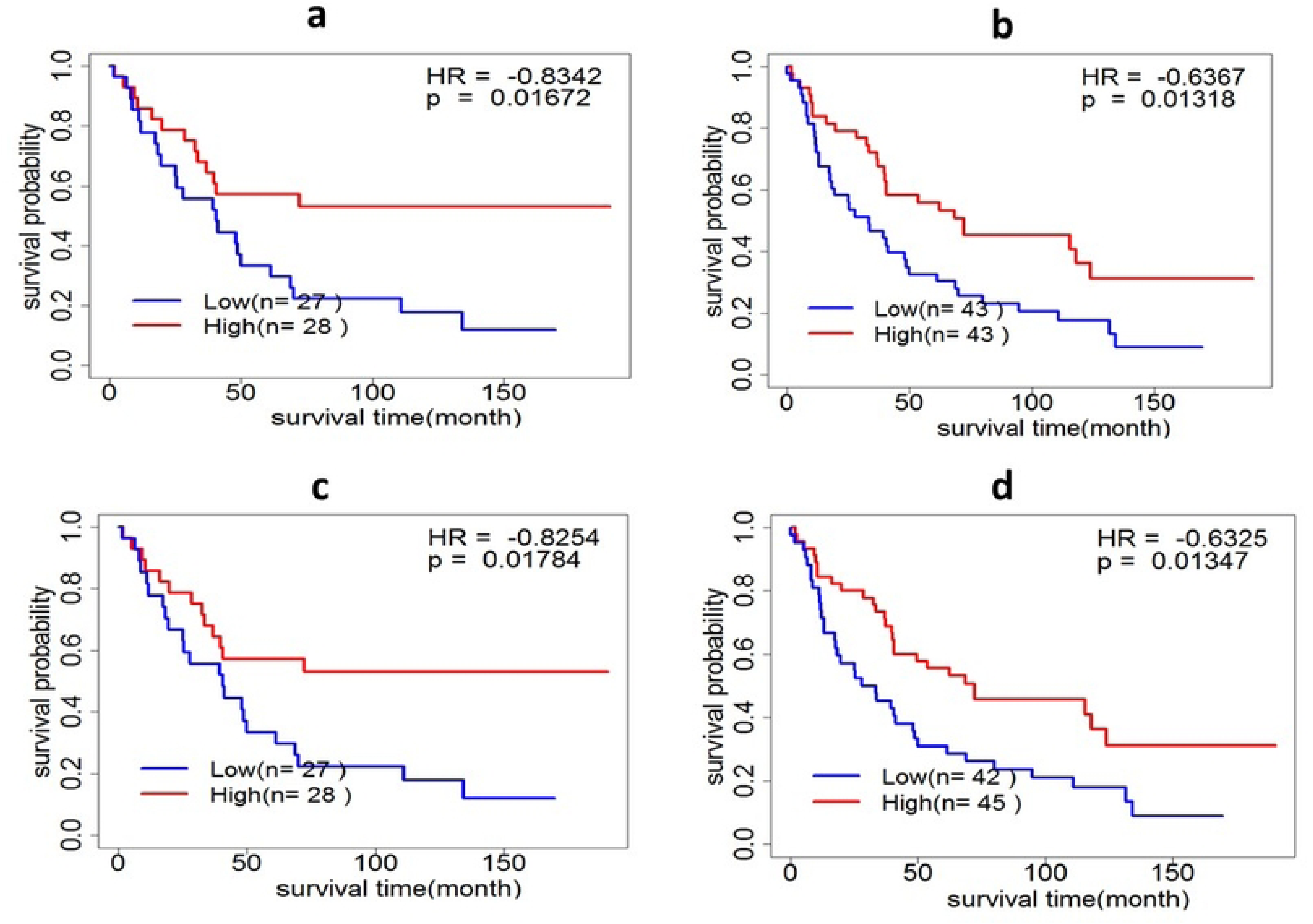
Cox proportional hazard survival analysis of NSCLC patients in the GSE37745 cohort GSE37745 consists of 196 lung cancer patients, of which 106 patients had ADC, 66 had SCC, and 24 had LCC. OS analysis was conducted for 106 ADC patients who died in stage 1 (**a**) and all stages (**b**) via a univariable Cox proportional hazard model and a multivariate Cox proportional hazard model adjusted for age and sex in stage I (**c**) and all stages (**d**).

#### 3.5.2. Survival analysis of patients with multiple cell types in death status using TS-signature

Overall survival analysis of the patients with ADC and ADC/SCC was conducted using our TS-signature. The results show that the high-expression patients had significantly higher survival probability than the low-expression patients with p=0.000349 (Fig.5a), 0.0027 (Fig.5b) in the univariable model and p-value = 0.00413 in the multivariate model adjusted for sex and age (Fig.5e) in the GSE8894 cohort and p-value = 0.01805 in the univariable model in the GSE3141 cohort (Fig.5). Compared with ADC, however, ADC/SCC clearly increased the death risk of patients (-0.8342 in the univariable model (Fig.5b), -0.7801 in the multivariate model in the GSE8894 cohort (Fig.5e), and -0.8798 in the GSE3141 cohort (Fig.5f).

In the GSE30219 cohort, we performed Cox proportional hazard survival analysis of 293 patients with ADC, SCC or other cell types using univariable and multivariate models. For the multivariate models adjusted for the covariates age, sex and/or tumor stages *t*, *n*, and *m*, the TS-signature did not change the death risk of patients in the multivariate model adjusted for age, sex, and tumor stage n (Fig.9d); however, it significantly decreased the death risk of patients with p-value=7.98e-08 for HR = -0.7395 ((Fig.9a) in the univariable model, and p-value = 8e-05 for HR = -0.642 in the multivariate model adjusted for age and sex (Fig.9b), and p-value= 0.0212 for HR = -0.4046 (Fig.9c) in model adjusted for age, sex, and tumor stage *t*, and p-value = 4e-05 for

**Figure 9.**
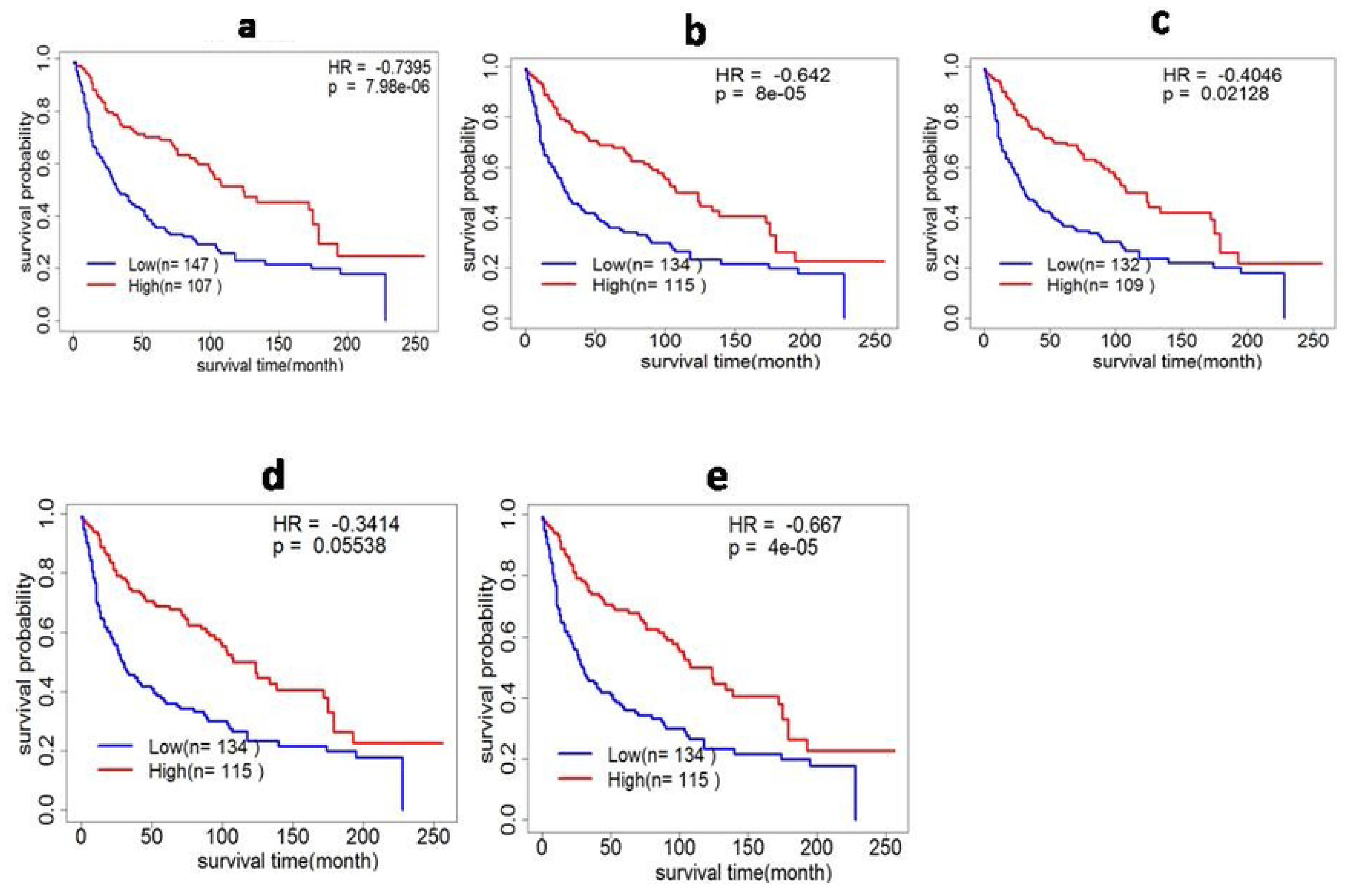
Cox proportional hazard survival analysis of the death status of patients with multiple cell types (ADC, SCC, LCC) in the GSE30219 cohort OS analysis was conducted for ADC, SCC or LCC patients who died via a univariable Cox proportional hazard model (**a**), multivariate Cox proportional hazard Model 1 (**b**), multivariate Model 2 adjusted for age, sex, and stage t (tumor size) (**c**), multivariate Model 3 adjusted for age, sex, and stage n (node) (**d**), and multivariate Model 4 adjusted for age, sex, and stage m (metastasis) (**e**).

**Figure 10.**
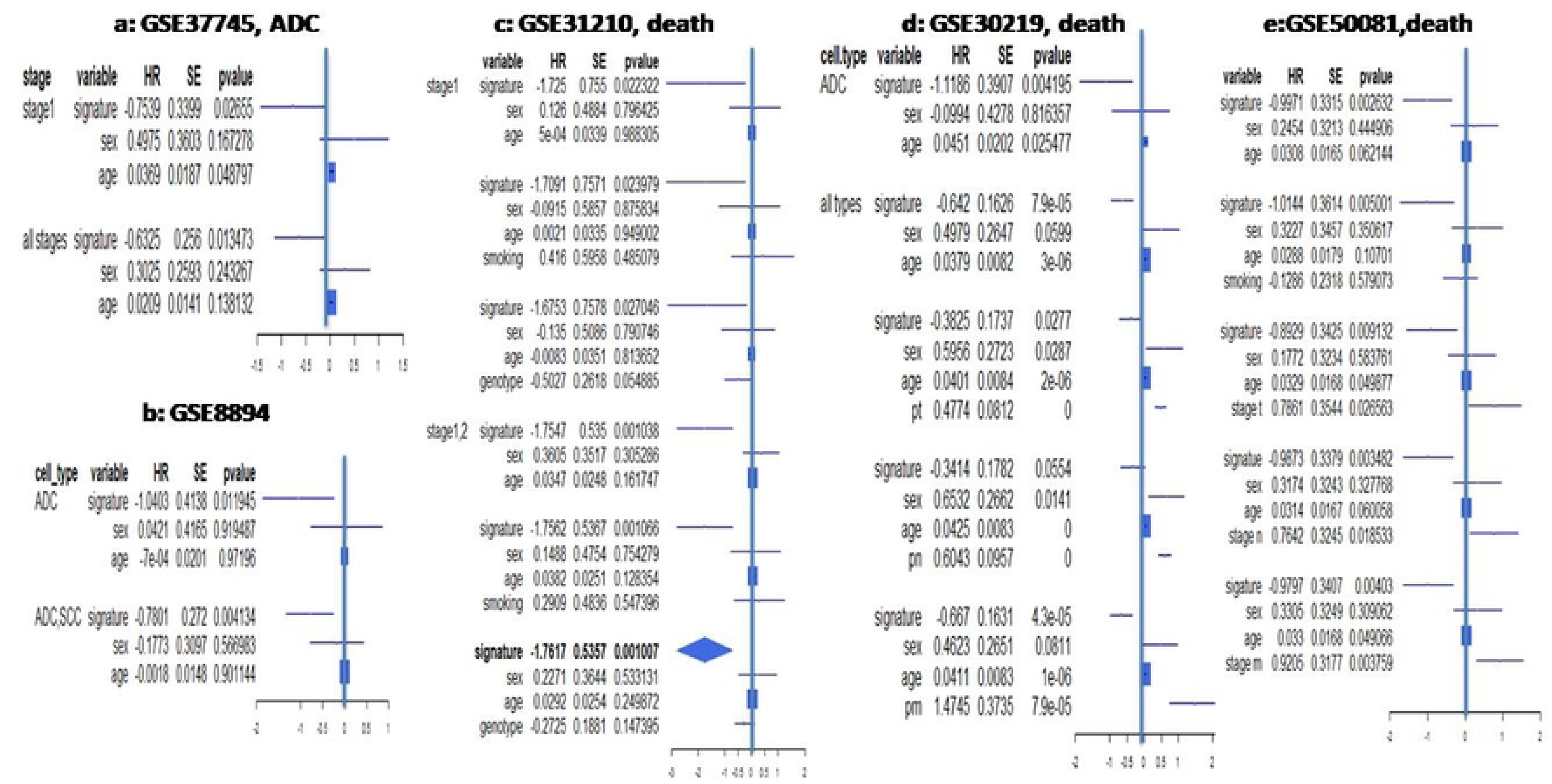
Forest plots for the results of multivariate Cox proportional hazard survival analysis of NSCLC patients The results of multivariate Cox proportional hazard survival analysis of ADC patients with death status in stages 1 and 2 in the GSE37745 cohort (a), ADC patients with death status in the GSE8894 cohort (b), ADC patients with death status in stages 1 and 2 in the GSE31210 cohort (c), ADC patients with death status in the GSE30219 cohort (d), and ADC patients with death status in the GSE50081 cohort (e).

HR = -0.667 (Fig.9c) in model adjusted for age, sex and tumor stage *m*. These suggest that, the high TS-signature expression patients had significantly higher survival probability than the low TS-signature expression patients. Compared with the multivariate model adjusted for age and sex (Fig.9b), we found that tumor stages *t*, *n*, and *m* significantly increased the death risk of patients (Fig.10d). Figure 10d also shows that tumor *stage* t had an HR = 0.4774 with p-value =4.096e-09, tumor stage *n* had an HR = 0.604 with p-value =2.754e-10, and tumor stage *m* had an HR =1.475 with p-value = 7.878e-05, suggesting that tumor stages *t*, *n*, and *m* were strongly positively associated with the death risk of patients with ADC, SCC or other types.

In the GSE42127 cohort, no significant differences between high- and low-expression patients with ADC/SCC in death status in stage1 or all stages were found in univariable and multivariate models (data not shown), indicating that the TS-signature does not respond to SCC. Tang et al. also reported that their signature did not predict the survival outcomes of patients with SCC in the GSE42127 cohort[5]. The covariates age, sex, and ACT (adjuvant chemotherapy) had no effect on the risk of death in stage 1 or all-stage patients with cell type ADC or SCC (Fig.S10).

#### 3.5.3. Survival analysis of ADC patients in relapse or recurrence status using TS signature

For patients with NSCLC, recurrence has a high risk of death, even after initial treatment with curative intent. Therefore, it is necessary to show ability of our TS-signature to evaluate or predict recurrence or relapse of patients. For recurrence in GSE50081, the patients with high expression of the TS-signature also had a much higher probability (70% after 8 months) of reducing recurrence risk than those with low expression (50% after 8 months). The patients with high expression of the TS-signature had hazard risk (HR) = -0.8895 with p-value = 0.01123 in the univariable model (Fig.11a) and HR= -0.9412 with p-value =0.01259 in multivariate model 1 adjusted for age and sex (Fig.11b) and model 2 adjusted for age, sex, and smoking (Fig.11c), HR= -0.7889 with p-value = 0.04652 in model 3 adjusted for age, sex, stage *t* (Fig.11d), HR= -0.8492 with p-value = 0.02872 in model 4 adjusted for age, sex, and stage *n* (Fig.11e), and HR= -0.8331 with p-value =0.03311 in model 5 adjusted for age, sex, and stage m (Fig.11f). In all multivariate models, the covariates age, sex, and smoking were not associated with relapse risk (Fig.S10A). However, tumor stages *n* and *m* significantly increased the relapse risk of patients with p-value < 0.01 (Fig.S10A).

**Figure 11.**
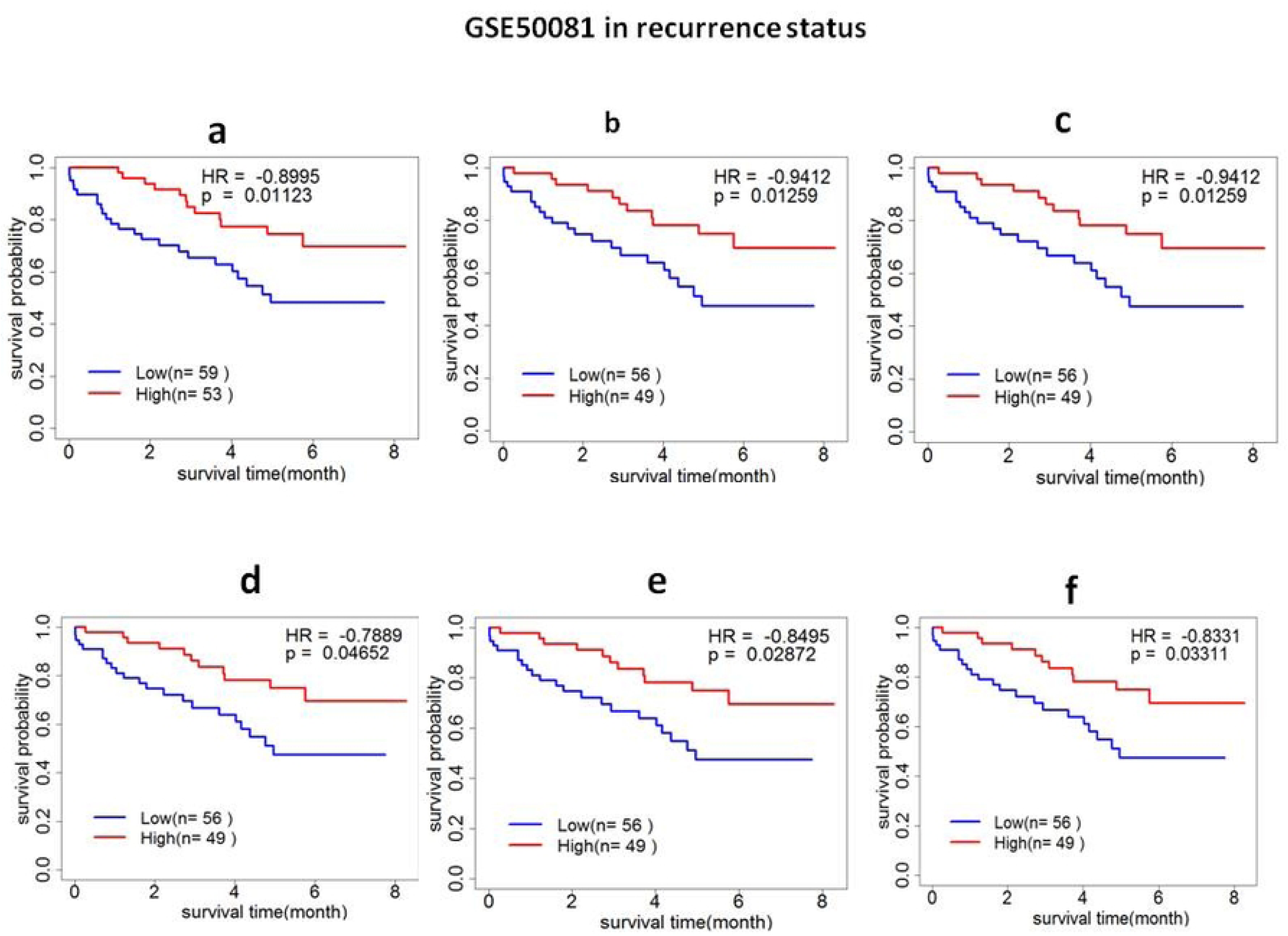
Cox proportional hazard survival analysis of NSCLC patients with recurrence status at all tumor stages in cohort GSE50081 Recurrence-free survival(RFS) analysis of ADC patients in cohort GSE50081, was conducted with univariable Cox proportional hazard model (**a**), multivariate Cox proportional hazard model1 adjusting for age and sex (**b**), multivariate Cox proportional hazard model2 adjusted for age, sex, and smoking (**c**), multivariate Cox proportional hazard model3 adjusted for age, sex, and stage t (tumor size) (**d**), multivariate Cox proportional hazard model 4 adjusted for age, sex, and stage n(node) (**e**), multivariate Cox proportional hazard model5adjusted for age, sex, and stage m(metastasis) (**f**).

GSE31210 also contains relapse data and censor time data from censors 1 and 2. We performed survival analysis of the 204 ADC patients with tumor stages 1-2 and time in censor1 via a univariable model (Fig.S7a) and multivariate model1 adjusted for age and sex (Fig.S7b); model 2 adjusted for age, sex, and smoking (Fig.S7c); and model3 adjusted for age, sex, and genotype (Fig.S7d). In addition, we also performed Cox proportional hazard analysis of 168 patients with stage-1 tumors via a univariable model(Fig.S7e) and multivariate Model 1 adjusted for age and sex (Fig.S7f); model 2 adjusted for age, sex, and smoking (Fig.S7g); and model 3 adjusted for age, sex, and genotype (Fig.S7h). The results are shown in Figure S7. According to the univariable model, HR = -1.0493 with p-value =0.00099 (Fig.S7a), similar to death, the patients with high TS-signature expression had a significantly higher probability (80%) of reducing relapse than those with low TS-signature expression (55%). After adjusting for age, sex and smoking status, the TS-signature had a greater negative hazard risk (HR=-1.1276) with a much lower p-value (0.00033) (Fig.S7c). Compared with Figure S7b (model 1 adjusted for age and sex), high TS- signature expression significantly decreased the relapse risk of patients when covariate smoking was added. Compared with model 1 adjusted for age and sex (Fig.S7b), after adjusting for genotype, the TS-signature obviously decreased the relapse risk of patients (HR= -0.9395 with p-value of 0.00281; Fig.S7d). In multivariate models 1-3 for patients in stage1, age, and sex were not significantly associated with relapse risk (Fig.S10C). In multivariate models 1-2 in stages 1-2, no associations of sex or smoking with relapse risk were observed, but age was found to be significantly associated with relapse risk (p-value=0.015 in Fig.S10C). In multivariate model 3, in stages 1-2, genotype significantly reduced relapse risk (HR=-0.331371) with p-value of 0.0245 (Fig.S10C).

To verify the utility of the TS-signature in stage-1-2 patients, we performed a sub-survival analysis of only stage-1 patients. This subset consisted of 163 patients in GSE31210. A statistically significant separation was observed in the Kaplan‒Meier survival curves of the 120-month ADC-related survival between the low- and high-expression groups in the univariable model (Fig.S7e), multivariate model 1 (Fig.S7f), model 3 (Fig.S7g), and model 4 (Fig.S7h). HR and p-value in multivariate model 1 adjusted for age and sex (HR = -1.045 with p-value = 0.0122 in Fig.S7f), model2 adjusted for age, sex, and smoking (HR = -1.032 with p-value = 0.0123 in Fig.S7g), and model3 adjusted for age, sex, and genotyping (HR = -1.045 with p-value = 0.0116 in Fig.S7h) are very similar to those in univariable mode (HR = -1.049 with p-value = 0.0129 in Fig.S7e). In the stage1 and stage 1-2 subsets, no covariate was found to be significantly associated with relapse risk, but genotyping still had a significant negative association with relapse risk (HR= -0.3841 with p-value = 0.0405 in stage-1 and HR = -0.3314 with p-value=0.0245 in stages 1-2) (Fig.S10C).

Survival analysis was conducted on the basis of the univariable and multivariate models adjusted for age, sex, smoking status and genotype in stage-1 and stage-2 patients. The results show that in stages 1-2, the TS-signature significantly reduced the risk of patients experiencing relapse with p-value = 0.00355 for HR=-0.9631 in univariable mode (Fig.S8a) and p-value = 0.00283 for HR = -0.9891 in multivariate model 1 adjusted for age and sex (Fig.S8b), p-value = 0.0024 for HR = -1.0405 in model 2 adjusted for age, sex, and smoking (Fig.S8c) and p-value = 0.00198 for HR= -1.0315 in model 3 adjusted for age, sex, and genotyping (Fig.S8d). The probability of cancer-specific survival in 110 months was 60% for patients with high TS-signature expressions and 35% for those with low TS-signature expressions. The difference between them was significant. In stage 1, we conducted survival analysis with a univariable model and multivariate models adjusted for age, sex, smoking status, and genotype. We found that the TS-signature effectively predicted the survival outcomes of patients and reduced the risk of relapse. Like relapse-censor1, the hazard risk of relapse occurred in patients was decreased in the multivariate models adjusted for age, sex, smoking status, and genotyping (HR< -1.0771), but p-values slightly increased (Fig.S8e, 8f, 8g, 8h). Fig.S10D shows that sex, smoking, and genotyping did not significantly impact relapse risk, but age was significantly positively associated with relapse risk in the stage-1-2 patients. However, we did not find that all these covariates were significantly associated with relapse risk in stage-1 patients (Fig.S10D).

Cohort GSE30219 also had relapse data, and our survival analysis of patients with ADC, SCC or other types was conducted with 5 models: a univariable model; multivariate model 1 adjusted for age and sex; model 2 adjusted for age, sex, and stage t; model 3 adjusted for age, sex, and stage n; and model4 adjusted for age, sex, and stage m. The results are summarized in Figure S9. In all five models, high-expression patients had a significantly higher probability of reducing relapse risk than low-expression patients in various situations, demonstrating that the TS-signature is a highly efficient and stable biomarker for the precise prediction of patient survival outcomes in different situations. In the GSE30219 cohort, the TS-signature also works well for predicting the prognosis of patients with lung cancer. **Figure S10B** shows that the covariates age and sex did not impact the risk of patients experiencing relapse, but tumor size (stage t) and node (stage n) very significantly increased the relapse risk of patients (HR=0.541 with p-value = 0 for tumor stage t and HR=0.606 with p-value = 0 for tumor stage n). Covariate metastasis (tumor stage m) also significantly promoted the risk of relapse in patients (HR=1.0737 with p-value=3.839e-02).

### 3.6. Comparisons of TS signatures with existing signatures

Many gene signatures for survival prediction in NSCLC patients have been published. Here, we arbitrarily chose four gene signatures to compare our TS-signature. The four gene signatures were respectively defined by He and Zuo[28], Chen et al[10], Navab et al[12], and Huang et al[29]. For convenience, we respectively called these gene signatures as the He-Zuo signature, Chen signature, Navab signature, and Huang signature. To evaluate the four signatures, we carried out univariable Cox survival analysis and used prognostic models to predict the overall survival of NSCLC patients with ADC and calculated their AUC values in the cohorts GSE3141, GSE8894, GSE50081, and GSE30219 (the four cohorts are typical cohorts: GSE3141 did not have covariates and had 52 ADC patients; GSE8894 had recurrence clinical data; GSE50081 had both death and recurrence data with multivariate age, sex, smoking and patients consisting of tumor stages *n, t,* and *m*; and GSE30219 has all cell types and tumor stages *n, t,* and *m* data. The four microarray datasets were produced from the GPL570 platform. The results are shown in Figure S11-S14 and in Table 1. Figure S11 shows that He-Zuo signature is similar to a tumor suppressor gene and has a good survival prediction ability (significant difference in survival probability between high- and low-expression patients with p-value =0.0077 in GSE3141 (Fig.S11A, p-value =0.00414 in GSE50081 with recurrence status data (Fig.S11C), and p-value =0.0002 in GSE50081 with death status data (Fig.S11D), but it did not work well in GSE8894 (Fig.S11B) and GSE30219 (Fig.S11E-11F. Table 1 shows that He-Zuo signature had better performance with an AUC = 0.6351 in GSE3141 and 0.6577 in GSE50081 (death and recurrence), but it had poor performance with an AUC < 0.56 in GSE30219 and an AUC=0.4336 in GSE8894. Therefore, He-Zuo signature is not a robust signature. Chen signature had poor survival predictions in all four cohorts (Fig.S12A-12E), except for relapse status in GSE30219 with p=0.01489 (Fig.S12F). For relapse/recurrence status, Chen signature had better performance for survival prediction in GSE8894 and GSE30219 datasets (AUC = 0.6173 and 0.6811, respectively) but had poor performance in GSE3141 and GSE50081. Navab signature had poor performance in GSE3141, GSE8894, GSE50081 (recurrence status), and GSE30219 (relapse status) (Fig.S13A-C,13F) with AUCs < 0.6 (Table 3), but in GSE50081 (death status) and GSE30219 (death status), its performance was good, with AUCs >0.6, which is in agreement with the results of the survival analysis (Fig.13D,13E). The Huang signature had better performance with an AUC of 0.6424 in GSE30219 (relapse status), but its performance was not good in GSE8894, GSE30219 (death status), GSE3141, or GSE50081 (recurrence and death statuses). Survival analysis revealed that Huang signature did not work in all four cohorts (Fig.S14A-14F). Our TS signatures had better performance in GSE50081 cohort (death and recurrence statuses) with AUC > 0.61, good performance in GSE3141, GSE8894 and GSE30219 (death status), with AUC > 0.7, and very good performance in GSE30219 (relapse statuses) with AUC> 0.8. These findings are consistent with the results of the overall survival analyses in these cohorts.

**Table 1.** AUC values of the signatures for the prediction of Cox proportional hazard models in the four cohorts.

|  | GSE3141 | GSE8894 | GSE50081<br>(recurrence) | GSE50081<br>(death) | GSE30219<br>(death) | GSE30219<br>(relapse) |
| --- | --- | --- | --- | --- | --- | --- |
| **He-Zuo Signature** | 0.6875 | 0.4336 | 0.6351 | 0.6577 | 0.5592 | 0.5356 |
| **Chen-Signature** | 0.5200 | 0.6173 | 0.5449 | 0.5256 | 0.5994 | 0.6811 |
| **Navab-Signature** | 0.5673 | 0.5604 | 0.5527 | 0.6096 | 0.6723 | 0.5931 |
| **Huang-Signature** | 0.5633 | 0.6040 | 0.5154 | 0.496 | 0.6069 | 0.6424 |
| **TS-signature** | 0.7424 | 0.7033 | 0.6295 | 0.6173 | 0.7165 | 0.8133 |

## 4. Discussion

Zhang et al[17] screened these 26 tumor suppressor genes and used S-score and N-score analyses characterized them. They used gene function analysis to reveal that these TS genes mostly work in the regulation of RNA transcriptions, protein translations, and transportations and are strongly correlated with the histone methylation genes and found that these 26 TS genes play important biological roles in tumor development. Here, we used dynamical differential expressions to show differential expression stage-course of the 26 TS genes between normal and tumor tissues in Taiwan female nonsmoking cohort[1]. As we expected, in the early stages of tumors, the 26 TS genes had weak expression differences between normal and tumor tissues but when tumor became late stages, almost all of the TS genes were highly expressed in normal samples but repressed in tumor tissues, suggesting that differential expressions of the TS genes can be used to identify developmental stages of tumors in ADC patients. In LMU Munich cohort (GSE103512), expressions of the TS genes in tumor cells K-cells, R-cells, P-cell, and NSCLC-coex-paths are similar to their expressions at stage 1A (Fig.1A). In Origene Technologies cohort (GSE40275), differential expressions of the TS genes between normal and tumor tissues are similar to stage 1B (Fig.1B). Therefore, the patient tumors in GSE103512 cohort can be inferred to be stage 1A and in GSE40275 cohort, development of patient tumors may be evaluated to be at stage 1A or 1B. Using microarray data GSE19804, we studied dynamical expression correlations of the TS genes with oncogenes through tumor stages and with genes associated with PD-1. The results demonstrate that at the early stages of tumors the TS genes were not obviously negatively correlated with oncogenes and tumor genes, however, when tumor developed to the late stages, there were strongly and negative correlations between the TS genes and oncogenes (or tumor genes), suggesting that developmental stages of patient tumors can also be diagnosed by using correlation of the TS genes with oncogenes. In addition, interestingly, maftool analysis of TCGA WES data showed that the 26 TS genes had very lower somatic mutation ratio than the oncogenes.

To confirm that our TS genes are every useful in therapeutic relevance, prediction and prognosis, we applied them to 7 different clinical and gene expression datasets. Unlike the risk score [4,12,15,30,31,32,33], we use negative log(p-values) (Eq.(1) and Eq.(2)) as weights to construct scores of the TS-signature across patients. Our TS-signature scores are utilized to perform univariable and multivariate Cox proportional hazard survival analyses of the ADC patients in death and recurrence/relapse statuses in the 7 cohorts. The results obtained from the 7 cohorts showed that the TS signature had high survival and recurrence predictions of ADC patients in NSCLC cancer. This is because the weights reflect contributions of genes to signature-scores, the TS-genes with lower p-values in single-gene survival analysis would have larger contributions to signature-scores while those with larger p-values would have small contributions. So, even though these TS genes have different expressions in different patients and in different cohorts, the signature-scores are mainly determined by most TS genes whose expressions are accordant in cancer patients. If one uses averages of expression values over 26 TS genes (weight=1/26) as signature-scores, each TS-gene has equal contribution to signature-score of each patient. Then the TS-genes whose expressions are highly associated with survival or recurrence status of patients cannot play main role in prognosis prediction of patients and the survival analysis would show very inconsistent results in different cohorts. In overall survival analysis, most studies use the median of the estimated risk score as the cutoff to classify patients into high- and low-risk groups[5,11,22,23,34,35,36]. In statistical theory, the median of the estimated risk score as the cutoff is equivalent to a z-score of zero in a standard normal distribution. A z-score of zero is highly uncertain because of noise, leading to inconsistent results across studies. In our method, we filtered out patients with -0.2 ≤ z-score ≤ 0.2, and high expression was defined as a z-score > 0.2, whereas low expression was defined as a z -score < -0.2. In other words, the standard normal distribution is a stable distribution and does not vary across cohorts or data. In addition, the 26 TS genes have stable and accordant properties in differential expressions in development of tumors. Therefore, our TS signature had robust predictions of death or recurrence/relapse risk. The results of the overall and recurrence-free survival analyses support this conclusion. Thus, the TS signature can be broadly applied to various cohorts, and the method for screening TS genes as signatures for survival analysis can be generalized to other types of lung cancer and other cancers.

## 5. Conclusions

In the early stages of tumors, the 26 TS genes had weak expression differences between normal and tumor tissues but in late stages, the TS genes were highly expressed in normal samples but repressed in tumor samples, implicating that differential expressions of the TS genes can be used to identify developmental stages of tumors in ADC patients.

Developmental stages of patient tumors can be diagnosed by using correlation of the TS genes with oncogenes or tumor genes.

The TS signature had high accurate prediction of prognosis in ADC patients and significantly reduced death or recurrence risk of ADC patients and the high TS-signature expression patients had significantly high survival probabilities than the low TS-signature expression patients.

Therefore, our TS signature can be used as a strong biomarker for detecting, predicting, and diagnosing cancer patients, especially, stage-early ADC patients.

## Abbreviations

NSCLC: Non-small cell lung cancer.
ADC: Adenocarcinoma.
SCLC: Small lung cancer.
SCC: Squamous cell carcinoma.
TS: Tumor suppressor.
ROC: Receiver operating characteristic
TPR: True positive rate.
FPR: False positive rate.
AUC: Area under the ROC curve.

## Data availability statement

The GSE3141, GSE8894, GSE68465, GSE30219, GSE50081, GSE37745, GSE42127, and GSE31210 datasets used in this study are public and can be downloaded from GEO (https://www.ncbi.nlm.nih.gov/geo/).

## Author contributions

**Man Jiang:** data collection, making figures and tables, writing-original draft preparation, funding acquisition, writing-reviewing, and investigation. **Yuan-De Tan:** conceptualization, writing-original draft preparation, visualization, investigation, writing-reviewing, visualization, investigation, methodology, statistical analysis, wrote software.

## Acknowledgements

We are grateful for supports of the Shangdong Provincial Health Technology Development Plan and the Shandong Natural Science Foundation.

